# Imaging erythrocyte sedimentation in whole blood

**DOI:** 10.1101/2021.08.31.458375

**Authors:** Alexis Darras, Hans Georg Breunig, Thomas John, Renping Zhao, Johannes Koch, Carsten Kummerow, Karsten König, Christian Wagner, Lars Kaestner

**Affiliations:** Experimental Physics, Saarland University, Saarbruecken, Germany; Biophotonics and Laser Technology, Saarland University, Saarbruecken, Germany; Department of Biophysics, Center for Integrative Physiology and Molecular Medicine, School of Medicine, Saarland University, Homburg, Germany; JenLab GmbH, Berlin, Germany; Department of Physics and Materials Science, University of Luxembourg, Luxembourg City, Luxembourg; Theoretical Medicine and Biosciences, Saarland University, Homburg, Germany

**Keywords:** red cells, erythrocyte sedimentation rate (ESR), mesoscopic microscopy, 2-photon microscopy, light-sheet microscopy, colloidal gel

## Abstract

The erythrocyte sedimentation rate (ESR) is one of the oldest medical diagnostic tools. However, currently there is some debate on the structure formed by the cells during the sedimentation process. While the conventional view is that erythrocytes sediment as separate aggregates, others have suggested that they form a percolating gel, similar to other colloidal suspensions. A direct probing of the structures formed by erythrocytes in blood at stasis is then required to settle these discrepancies. Here, we report observations performed with three different optical imaging techniques: direct light transmission through thin samples, two-photon microscopy and light-sheet microscopy. All techniques revealed a dynamic structure of a channeling gel but with differences in the resolved details. A quantitative analysis of the erythrocyte related processes and interactions during the sedimentation need a further refinement of the experimental set-ups.

## 1. Introduction

The erythrocyte sedimentation rate (ESR) is one of the oldest diagnostic methods. Already in ancient times it was known that the sedimentation of the red part of the blood can be very different (Kushner, 1988). In fact, it was known even before the thermometer was invented that many diseases cause a higher ESR (Bernhardt and Hianik, 2000), which as we know today is also related to increased body temperature.

At the end of the19th century, the ESR was developed into a diagnostic tool very close to how it is used nowadays (Biernacki, 1894; Westergreen, 1921). It was recognized as an non-specific measure of inflammation with all the advantages and disadvantages of an unspecific test.

The primary explanation of the process goes back to the seminal work of Robin Fåhraeus (Fahraeus, 1929) and is based on the effect that erythrocytes form aggregates phenomenologically described as the formation of ‘stack of coins’, also known as “rouleaux”. There has been a long dispute whether this is caused by depletion or specific binding and probably both mechanisms contribute to the effect (Baskurt et al., 2012; Flormann et al., 2017). Medically, there are two effects involved: (i) the increase of plasma proteins associated with inflammation. This is the case for fibrinogen. Although also C-reactive protein (CRP) is increased in inflammation, it does not contribute to increased ESR (Flormann et al., 2015). In contrast, (ii) also the erythrocyte shape contributes to the ESR. Irregular cell shapes, like sickle cells (Jan et al., 1981; Lawrence and Fabry, 1986) or acanthocytes (Salt et al., 1960) decrease the ESR. Although there is no lower limit established as a pathological limit, a low ESR was proposed as a biomarker for diseases belonging to the neuroacanthocytosis syndrome (NAS) (Darras et al., 2021; Rabe et al., 2021).

The conventional view of erythrocytes sedimentation is that erythrocytes sediment as separate aggregates. This assumption suits the observation of a higher sedimentation rate for higher blood plasma protein concentrations, inducing a higher aggregation force between the cells. Indeed, in such a situation, the bigger the aggregates are, the faster they sediment. However, it is at odds with the existence of a sharp phase transition between plasma and erythrocyte suspension that is clearly visible in all phases of the sedimentation process. This could be explained by the fact that erythrocytes aggregate into a percolating gel that collapses during the sedimentation process (Darras et al., 2021), as it is the case for other colloidal suspensions. Regarding blood, the position of the sharp transition interface can oscillate in time (Tuchin et al., 2002). The evolution of the suspension’s electrical conductivity over time also supports the idea that the structure evolves into a gel containing some plasma channels (Pribush et al., 2010). However, classical colloidal gels tend to sediment slower when the attraction between the suspended particles increases (Gopalakrishnan et al., 2006), then supporting the conventional view. To settle these discrepancies and better understand the fundamental mechanisms of erythrocyte sedimentation, it is necessary to have a closer mesoscopic or microscopic view on the sedimentation process. Due to the high optical density of erythrocytes in whole blood, this microscopic task is a challenge. Furthermore, we have to match the need to image perpendicularly to the gravitational force, which excludes classical upright and inverted microscopes.

Here, we compare three different approaches: (i) mesoscopic transmission microscopy with a microscope put on its side, (ii) two-photon microscopy with a special device with an objective mounted on a hinched bracket designed for *in vivo* investigations of human skin (König, 2018) and (iii) light-sheet microscopy, which intrinsically has the right geometry.

## 2. Methods

We attempted to highlight the structure adopted by erythrocytes during sedimentation with three different imaging techniques. The first one is a simple transmission of blue light through a thin sample. The two others are advanced but well-established techniques, namely two-photon microscopy and light-sheet microscopy. This section details the sample preparations and the basic principles of each technique.

Blood samples were collected as previously described (Abay et al., 2019). Blood was exclusively used from healthy volunteers with an informed consent, according to the declaration of Helsinki and the approval by the ethics committee ‘Ärztekammer des Saarlandes’, ethics votum 51/18. Nine ml of blood were collected in EDTA-containing tubes (Sarstedt, Nümbrecht, Germany). If indicated, a fluorescent dye was added and then the blood directly transferred into the measurement containers. All measurements were performed within six hours after withdrawal.

To assess the effect of the container’s geometry on the ESR, we performed classic ESR measurements with the same healthy sample with a controlled hematocrit of 45%. The height of cell-free plasma layer on top of erythrocytes after 1h are respectively 9.3 ± 0.3mm, 7.5 ± 0.3 mm and 9.9 ± 0.3 mm for the transmission, two-photons and light-sheet microscopy. Sedimentation velocity of the erythrocytes in the various containers then have the same order of magnitude.

### 2.1 Mesoscopic blue-light transmission imaging

We first tried to image the structure of erythrocytes during sedimentation in a thin container via direct blue-light transmission. We chose blue light because this provides the highest contrast due to the Soret band in the absorption spectrum of hemoglobin (Kaestner et al., 2006).

We prepared containers with two microscope glass slides (VWR, Radnor, Pennsylvania, USA), separated by two bands of paraffin, 150±50 μm thick (see Figure 1A for a scheme and Figure 1B for a picture of the container). To ensure homogenous wetting, both plates were first washed with isopropanol and distilled water, then dried with clean compressed air. The paraffin layer was made by cutting a few Parafilm^®^ M (Carl Roth GmbH, Karlsruhe, Germany) bands, placing them on one of the glass slides and then heating the system briefly to 70°C on a heating plate (RCT basic, IKA-Werke GmbH, Staufen, Germany). Once the paraffin became malleable, the second glass plate was placed on top and the whole system was gently pressed together by hand.

**Figure 1:**
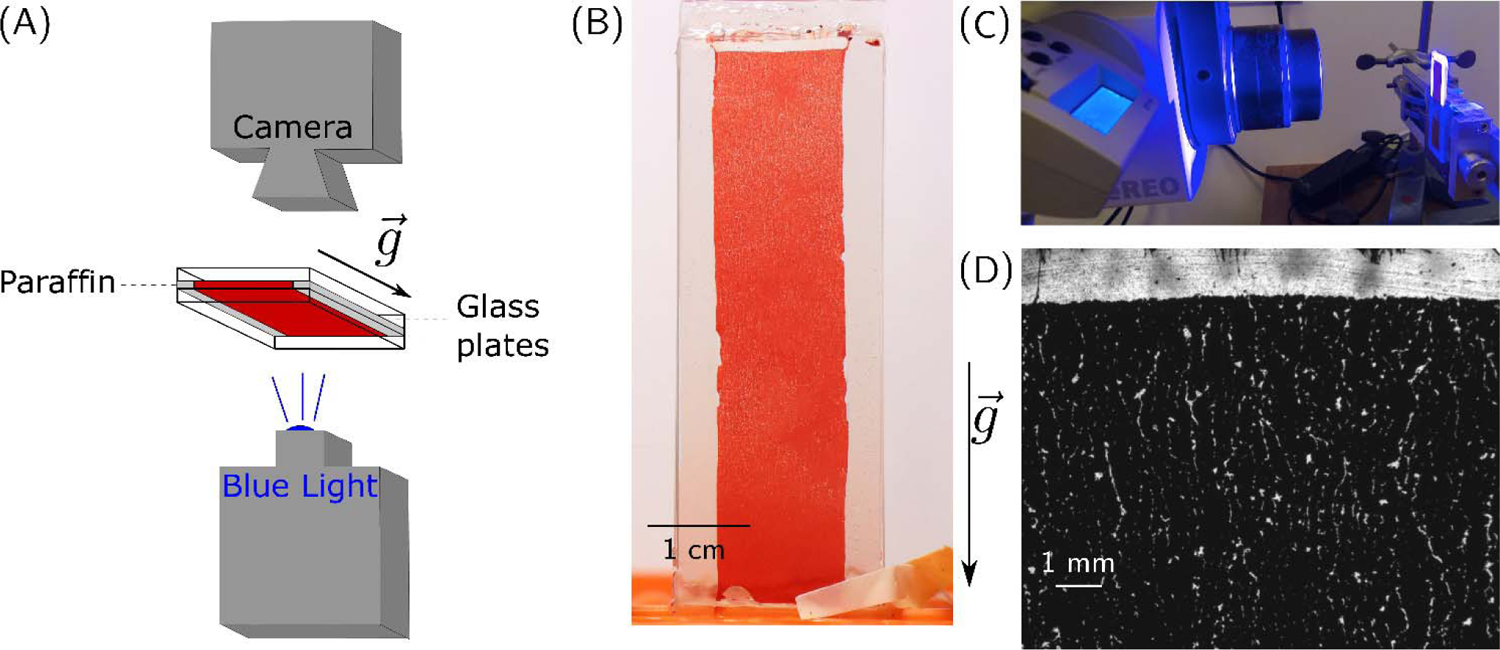
Mesoscopic blue-light transmission observation. (A) Scheme of the microscopy set-up. (B) Picture of the whole sample, as obtained from a regular camera with macroscopic objective. The picture has been obtained after 90 min sedimentation time. (C) Picture of the microscopy set-up. (D) Microscopic picture obtained with blue light transmission after 30 min sedimentation time.

The blood sample was completely drawn into the space between the plates by capillary forces. The bottom and top openings were then sealed with petroleum jelly (KORASILON-Paste, Kurt Obermeier GmbH, Bad Berleburg, Germany).

For high-resolution pictures of the observed cell-free areas, we illuminated the samples with blue light (High-power LED SOLIS-415C, Thorlabs, Newton, NJ, USA). Transmission images were then obtained with a microscope whose observation axis was horizontal. Figure 1A shows the scheme of the set-up and Figure 1C the corresponding set-up of the Stereomicroscope (Zeiss, Jena, Germany) with an 8x zoom objective.

### 2.2 Two-photon microscopy

Two-photon microscopy is a technique which is applied whenever a high penetration depth into tissue is required, e.g., when imaging skin layers *in vivo* (<u>König and Riemann, 2003</u>; Breunig et al., 2012; König et al., 2015; König, 2018), brain structures (Denk et al., 1995; Cartarozzi et al., 2018) or cardiac tissue (Kaestner and Lipp, 2007; Hammer et al., 2014). Here, we used the MTPflex CARS tomograph (JenLab GmbH, Berlin, Germany) with a flexible arm, originally designed to assess skin lesions by *in vivo* imaging (Weinigel et al., 2015).

A 30 mM stock suspension of free-acid fluorescein (Sigma-Aldrich, Missouri, USA) in phosphate buffered saline (PBS, Gibco, New-York, USA) was prepared. For the observations, erythrocytes and plasma were first separated by centrifugation. Subsequently, 10 μL of the fluorescein 30 mM stock suspension were added to 1 mL of plasma, thereby creating a 300 μM concentration of fluorescein. The dyed plasma was then mixed again with packed erythrocytes to reach a final hematocrit of 45%. When macroscopically compared with erythrocytes resuspended in pure plasma, the dyed suspension did not present any significant modification of the ESR.

We used spectroscopy cuvettes with a square cross-section of 3×3 mm^2^ and 20 mm high (Art. No. 101-015-40, Hellma Analytics, Müllheim, Germany). The cuvettes were filled with 180 μL suspension by the means of a micropipette. They were then sealed by a cover glass slide, on which a thin layer of petroleum jelly had been spread, and the cuvette was eventually turned upside-down for a better reproducibility of the top surface. For imaging, femtosecond laser pulses were focused into the sample and the focal position scanned across the region of interest. Fluorescence signals were pixel wise collected in reflection geometry to generate the two photon images. The mean laser power was approximately 38 mW, the laser wavelength was set to 800 nm. A low NA objective (Fluar 5x/0.25NA, Zeiss, Germany) was employed to enable a large field-of-view. Figure 2A and B show the tomograph and the sample setup, respectively.

**Figure 2:**
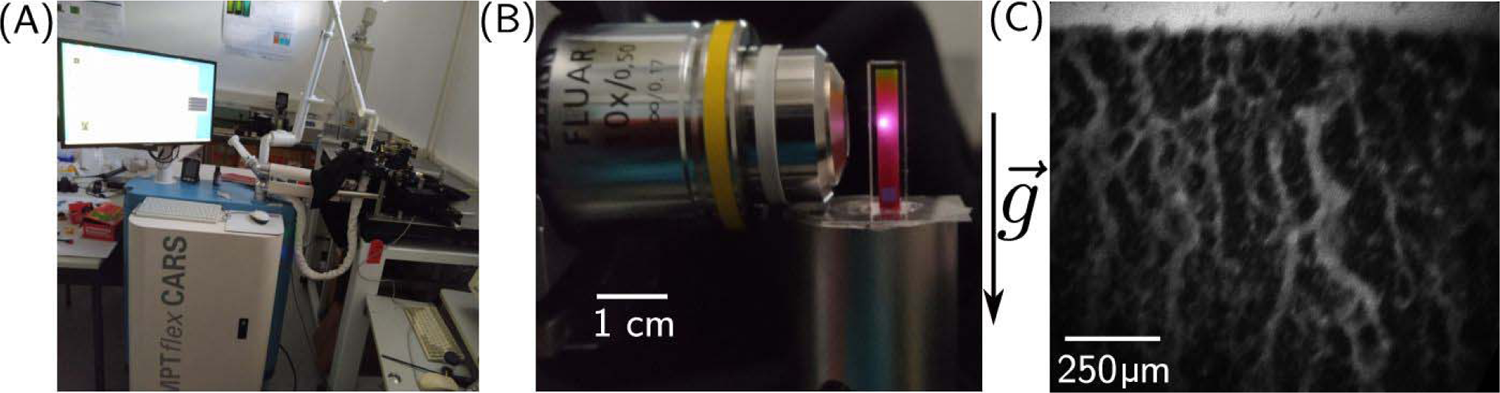
Two-photons microscopy. (A) Photograph of the whole MTP flex CARS tomograph, used for two-photon imaging. (B) Picture of a sample and the microscopy set-up. (Picture taken for illustration purpose, the laser isn’t focused at the observation plane, which is the surface of the sample.) (C) Picture obtained of the erythrocytes configuration after 30 min sedimentation time

### 2.3 Light-sheet microscopy

Light sheet microscopy is based on separate light paths for sample excitation and image acquisition (Huisken et al., 2004). It has in common with the two-photon microscopy that fluorophores are excited only in the layers where image generation occurs. This is in contrast to confocal microscopy and other sectioning microscopy methods (Lipp and Kaestner, 2006) and has the advantage to avoid unnecessary photobleaching of the samples.

Finally, we prepared a 10 mM stock solution of Atto 647 carboxyl (Atto-Tec, Siegen, Germany) in PBS. For the observations, erythrocytes and plasma were first separated by centrifugation. Subsequently, 1 μL of the dye solution was added to 1 mL of plasma, thereby creating a 10 μM concentration of Atto647. The stained plasma was then mixed again with packed erythrocytes to reach a final hematocrit of 45%.

To prepare the sample, 60 μL of the stained erythrocyte suspension was then placed in a fluorinated ethylene propylene (FEP)-tube (KAP 101.653, Techlab, Braunschweig, Germany) with inner diameter of 1.6 mm. The bottom of the tube was sealed with petroleum jelly. The sample was then placed inside the light-sheet microscope chamber (Z1, Zeiss, Jena, Germany) and illuminated with a laser sheet, as illustrated schematically in Figure 3A and in the actual device Figure 3B.

**Figure 3:**
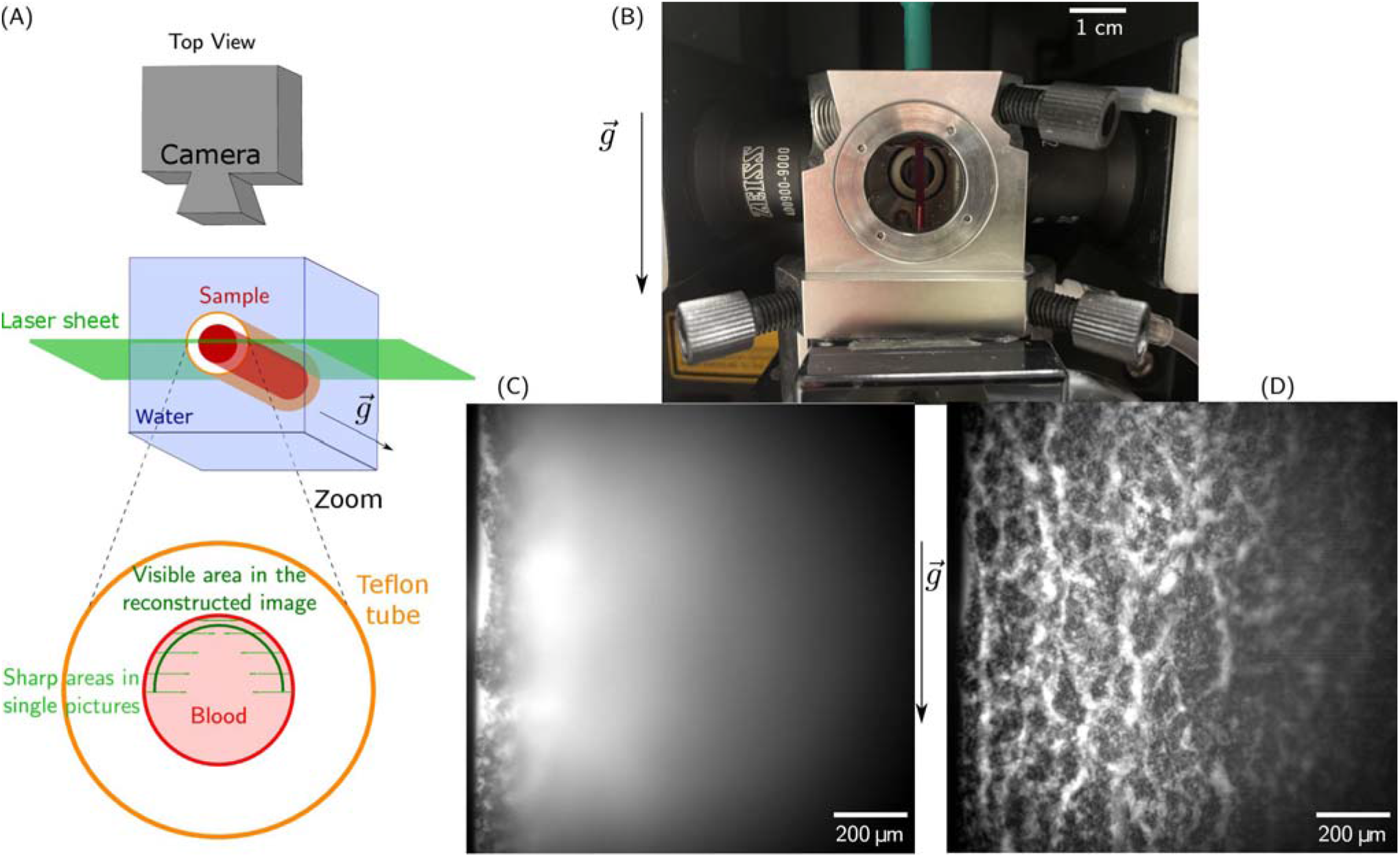
Light-sheet microscopy. (A) Scheme of the setup. The position of the taken pictures and the location of the reconstructed are also represented. (B) Picture of the set-up. The light-sheet lenses are on the left and right side, the recording camera is behind the sample. (C) Raw picture as obtained during the experiment. (D) Reconstructed picture obtained after postprocessing of a *z*-stack. This stack was obtained after 90 min sedimentation time of the sample.

A *z*-stack, recorded with a 20x objective and a 0.4x post reduction, was performed by modifying the position of the sample relative to the light sheet in steps of 5 μm, with a total of 74 slices, in approximately 2s. Given the high-volume fraction of the erythrocytes, the region suitable for imaging is actually limited to approximately 100 μm from the sample border. However, one can then reconstruct a complete picture by averaging the areas in which structures appear sharp from the different pictures of the *z*-stack.

For the picture reconstruction, a dedicated Matlab (MathWorks, Natick, Massachusetts, USA) routine was implemented. An inhomogeneous background level was identified for each picture through morphological opening using a circle with a radius of 60 μm as a structuring element. The background levels were then removed from the initial images and the resulting filtered images were averaged.

## 3. Results

### 3.1 Mesoscopic blue-light transmission imaging

Figure 1C and D show the setup with the blood sample in between the two coverslips separated by a distance of approximately 150 μm and a representative image obtained with the stereo microscope (8x zoom), respectively. Additionally, we present a movie of the sedimentation process in the supplemental material (Movie 1). The suspension is initially homogeneous, but mesoscopic channels appear in the container within 15 min. The observation of some winding cell-free channels is already possible in daylight by eye or with a macroscopic objective and a regular camera. However, the characteristic width of such channels is typically a few tens of micrometers. Since the thickness of the container itself was already approximately 150 μm, it is clear that only parts of the biggest cell-free channels were actually observable with this technique.

The images highlight the appearance of cell-free crack-like structures and indicate the potential for easily reproducible mesoscopic observations. It then qualitatively supports the hypothesis that erythrocytes organize into a percolating colloidal gel, and already leads to the observation of a characteristic distance of the order of a few hundreds of micrometers between two of the visible cracks. These first observations then imply that further observation techniques should cover an area of approximately 1 mm wide to capture a statistically relevant part of the sample organization. However, these pictures cannot resolve the actual size and properties of the channels, nor the detailed cell organization. Furthermore, the width of the samples is only reproducible with an accuracy of 33%, due to its handcrafting. This creates almost a ratio of two between the thickness of the thinnest (thickness of 102±20 μm) and the thickest containers (average thickness of 170±20 μm), as measured on microscopic images of the container sides. We then turned towards more sophisticated microscopy techniques, in order to improve the quality of the information obtained.

### 3.2 Sample staining concepts

While the transmission imaging works marker free with unstained samples, the following microscopy approaches require fluorescence staining. In an initial attempt we aimed to stain the erythrocytes. To be outside the main absorption of the hemoglobin (Kaestner et al., 2006) we looked for dyes that could be excited with a 640 nm laser or appropriate two-photon excitation wavelengths. Furthermore, we wanted the dye to be cytosolic in order to avoid any changes in the membrane structure and hence putative erythrocyte interaction properties. Therefore, we chose Cell Tracer deep red and Cell Tracker far red. Both dyes gave in principle nice erythrocyte staining as exemplified in Supplemental Figure S1A. However, the erythrocyte staining required the separation of the erythrocytes from the plasma (otherwise all dye is bound to albumin in the plasma) and a resuspension of cells after passing the staining procedure. We did not investigate the mechanism, but this staining procedure significantly altered the ESR (Supplemental Figure S1B and Supplemental Movie 2) and was therefore rejected. Instead, we aimed for a staining of the plasma and performed it with Atto 647 carboxyl and fluorescein as detailed above. When macroscopically compared with erythrocytes resuspended in pure plasma, the dyed suspension did not present any significant modification of the ESR as outlined in Supplemental Figure S2.

### 3.3 Two-photon microscopy

The two-photon microscopy is normally performed with high-NA objectives since the non-linear intensity dependence of the excitation requires very high intensity thereby providing intrinsic high-resolution sectioning capability with limited field of view. However, in order to maximize the imaging region a low NA objective was employed here. Figure 2C shows a picture of blood in a cuvette obtained by two-photon microscopy with a 5x/ NA 0.25 objective. The achievable resolutions for this objective were estimated to be 2 μm and 20 μm lateral and axially, respectively, while providing a maximum image region of approximately 2×2 mm^2^.The corresponding focal volume is then of 125 μm^3^.

The two-photon microscopy offers a real-time transverse view of the erythrocyte organization. Indeed, as estimated from classic ESR measurements, instantaneous sedimentation speeds are up to approximately 8 μm per seconds. The two-photon microscopy allows us to take one image in approximately 4s, meaning that the cells can then move up to ~30μm within one frame recording. Since we image a gel with a characteristic width of a few millimeter, this maximal displacement is around 1% of the total structure size, and is therefore negligible when compared to the characteristic size of the image and the observed structures.

This is also highlighted in the corresponding movie of the sedimentation process in the supplemental material (Movie 3). The images nicely resolve channels appearing in a continuous structure. However, with the low NA objective, the erythrocytes cannot be clearly resolved in the bulk, and the initial state of the gel is not resolved at all. Moreover, due to the high absorption of the emitted light by the sample, the focal position had to be set close to the lateral face of the gel. Its exact position is actually the position of the only plane where we can obtain a sharp image at physiological volume fractions. We then assume that it is at the lateral face, within a range equal to the focal depth (20 μm).. Eventually, the illumination profile follows a Gaussian intensity profile, making it difficult to analyze quantitatively a large area.

Subsuming the two-photon imaging approach, this technique offers a real time direct observation of the channels at the lateral face of the colloidal gel. However, the field of view and resolution of the imaging in our setting is limited, and the individual cells cannot be resolved.

### 3.4 Light-sheet microscopy

The light-sheet microscopy has a default configuration that fits the measurement geometry required for the erythrocyte sedimentation (cp. Figure 3A). However, the samples are usually embedded in a matrix (gel), which is then suspended in a physiological solution. This works well for individual cells (Schoppmeyer et al., 2018; Backes et al., 2018), embryos (Huisken et al., 2004) or even isolated sinoatrial nodes (Flügel et al., 2018). However, this kind of sample preparation would completely alter the erythrocyte sedimentation process, therefore we had to go down new routes. We looked for a transparent material with an optical refraction index similar to water, and we identified fluorinated ethylene propylene tubes as a suitable cylindrical geometry that is compatible with both the erythrocyte sedimentation in blood and the light-sheet microscope from sample holder to optical arrangements (Figure 3A).

As depicted in Figure 3C, the penetration depth of the light sheet is limited, but based on a *z* stack of such images, we managed to reconstruct an image representing a curved plane as outlined in Figure 3A. An example of such a reconstructed average picture is shown in Figure 3D. A movie based on such reconstructed images is provided in the supplemental material (Movie 4).

The raw pictures obtained from the light sheet microscopy do not provide a complete overview of the structure adopted by aggregated erythrocytes during their sedimentation, since the penetration depth was limited to approximately 100 μm by absorption and scattering (Figure 3C). A live observation of the entire sample is therefore not possible. However, combining these pictures afterwards allows to obtain a good estimation of the real-time process with a curved interface structure and a time resolution limited to the acquisition time required for one z-stack. For the Supplemental Movie 4, one scan of the sample was performed every 7.9s, with a step of 5 μm between each of the 74 steps. The last picture of the scan lies at the front interface of the tube. Each slice is taken in 105 ms.

Even though the obtained pictures do not represent an actual cross-section of the 3D structure adopted by the erythrocytes during their sedimentation, the resulting pictures have a higher resolution compared to the other techniques tested. Some single cells are even resolved in the final processed picture. Overall, this imaging technique clearly shows that erythrocytes aggregate into a percolating structure, i.e. a colloidal gel, containing dynamic plasma channels allowing this liquid to flow up to the gel surface. The required post-processing prevents a live observation of the whole structure, which might be a limitation for more complex experimental protocols.

## 4. Discussion

Table 1 shows a comparison of the various advantages and drawbacks of each imaging technique. The following subsections also sum up the main original observations obtained with each technique.

**Table 1:** Comparison of the imaging techniques.

|  | **Sample**<br>**Reproducibility** | **Live observation** | **Simultaneous**<br>**resolution of**<br>**erythrocytes and**<br>**plasma channels** | **Actual cross-**<br>**section of the**<br>**sample** |
| --- | --- | --- | --- | --- |
| **Blue light**<br>**mesoscopic**<br>**transmission**<br>**imaging** | variable thickness of<br>the sample due to<br>hardly reproducible<br>sample preparation | yes, acquisition<br>speed only limited<br>frame rate of<br>camera | no, mesoscopic<br>imaging, just the<br>high contrast<br>plasma channels<br>are visible | limited, as we<br>image a<br>projection of the<br>entire thickness<br>of the sample |
| **Two-photon**<br>**microscopy** | sampling in<br>standardized optical<br>cuvettes | yes, acquisition<br>speed limited by<br>the scanning<br>process | limited resolution<br>within a relevant<br>field of view;<br>potential for<br>improvements | well defined<br>cross section of<br>a plane in the<br>sample |
| **Light sheet**<br>**microscopy** | medium, the tube has<br>a roughly constant<br>diameter but sample<br>is prone to slight tube<br>bending | no, a single plane<br>image requires the<br>recording of an<br>entire stack; image<br>generation needs<br>stack processing | best resolution<br>among the<br>systems tested | calculated image<br>along the inner<br>surface of the<br>circular tube,<br>sectioning<br>occurs in<br>principle but<br>hard to specify |

### 4.1 Mesoscopic blue light transmission imaging

The unique feature of the transmission imaging concept compared to the other methods is the marker free character, i.e. the sedimentation process is completely undisturbed. However, this holds only true for the composition of the blood, whereas the container has probably the biggest influence of the three methods tested: the distance of the coverslips of approximately 150 μm results in ‘wall effects’ (Starrs et al., 2002) most probably in the entire volume. Nevertheless, we are able to demonstrate the channel formation and its dynamics and such support the concept of the erythrocyte sedimentation of whole blood as a process which can be described as a percolating gel.

### 4.2 Two-photon microscopy

Two-photon microscopy is the technique that theoretically should be ideally suited to image blood sedimentation. A sub-micrometer optical resolution and a tissue penetration depth of up to 1 mm (Denk et al., 1995) should allow imaging of erythrocytes in a cuvette, with a cellular resolution and at sufficient distance from the cuvette wall to prevent wall effects. However, this applies to ideal conditions using high magnification objectives with a high numerical aperture. Due to the nature of the process, we want to image a field of view with a width of the order of a millimeter, which made us choose a 5x objective (with a numerical aperture of 0.25). Such an objective is far away from the ideal conditions sketched above. Although the laser scanning process results in a temporal delay of the points/pixels of the image, the sedimentation process is slow enough to regard the two-photon image acquisition as quasi live recording.

Furthermore, the dye / laser wavelength combination (fluorescein / 800 nm) was selected for highest signals, which is perfectly fine considering the excitation process, because the 800 nm is outside the hemoglobin absorption peaks and the two-photon excitation (corresponding to approximately 400 nm) needs to happen outside the erythrocytes in the plasma. However, the emitted light around 520 nm is still absorbed to a considerable degree by the hemoglobin and this loss in signal could be minimized with a dye emitting around 700 nm, which in turn would also require a laser excitation wavelength around or above 1200 nm, which is not a standard in most two-photon microscopes. Additionally, the light scatter induced by the lipids of the erythrocytes is a parameter that should be considered in further approaches.

In spite of all these limitations, the resolution we get is highly improved compared to the mesoscopic approach described above and although individual cells could not be resolved, we get a more detailed impression of the hydrodynamic processes occurring during the gel percolation.

### 4.3 Light sheet microscopy

Also, for the light sheet microscopy we faced hardware limitations. The 20x objective was the one with the lowest magnification available on site and a 0.4x post reduction was required to achieve a field of view with a width of 1.2 mm. Nevertheless, the light sheet microscopy is very promising to extract a quantitative description of the percolating gel, if one can establish a reproducible observation protocol which does not require a live feedback to the observer. In our opinion, this technique then offers the best resolution possible, but does not show an actual linear cross-section of the structure. Nonetheless, the images obtained from the reconstruction of the z-stack shows a realistic structure when compared with the two-photon microscopic images. The description of the surface geometry can then be acceptably described by the provided pictures. However, this does not offer any guarantee that the structure in the bulk of the structure is similar, which is a possibly problematic outcome when investigating the detailed colloidal physics process. However, the surface properties of fluorinated ethylene propylene (teflon) might have advantageous properties compared to the glass surface of cuvettes. Certainly, the good match of refractive indices of FEP and water and the higher aperture of the water immersion lens (NA 1.0) yielded a good resolution of the sedimentation process.

## 5. Conclusion

All imaging methods tested undoubtedly proved that erythrocyte sedimentation occurs as the dynamic compaction of a colloidal gel with plasma channels. Namely, all pictures showed a cohesive and percolating structure of erythrocytes containing plasma channels. Such structure is what defines a colloidal gel. Moreover, the Supplementary Movies clearly show that the erythrocytes sediment with a continuous velocity field, and not discrete velocity values as expected for disjoint aggregates. The only sudden velocity variations are in the plasma channels, where the liquid is flowing upwards and can rip off and drag up some small erythrocyte aggregates from the cohesive structure. Thus, all methods were able to allow insights into the process of the percolating gel during erythrocyte sedimentation of whole blood, however the channel formation dynamics needs to be investigated in more detail in further studies. All methods have their advantages and disadvantages as outlined in Table 1, and therefore we cannot select one method as being best suited. A limitation of all methods is the limited penetration depth into the blood and therefore the imaged processes might be influenced by erythrocyte interactions with the respective container walls. In that respect, multi-photon imaging has the highest optimization potential for improvements by, e.g., the application of bathochromic shifted laser wavelength or use of adaptive optics.

## 7. Author Contributions

The authors contributed the following to the manuscript: Conceptualization, A.D. and L.K.; methodology, A.D., H.G.B., T.J., R.Z., J.K., C.K. and L.K.; software, A.D.; investigation, A.D., H.G.B., R.Z. and J.K.; resources, K.K. and C.W.; data curation, A.D. and L.K.; writing—original draft preparation, A.D. and L.K.; writing—review and editing, all authors; visualization, A.D. and L.K.; supervision, C.K., K.K. and C.W.; project administration, L.K.; funding acquisition, C.W. and L.K. All authors have read and agreed to the published version of the manuscript.

## 9. Acknowledgements

We acknowledge the support of the project by Prof. Markus Hoth (Biophysics, Saarland University).

