## Supplemental Movies' Legends for "Imaging erythrocyte sedimentation in whole blood"

**Supplemental Movie 1: Mesoscopic blue-light transmission observation.** (File ChannelsE4Hem45S0_20200909SpedUp50X.avi) A suspension of erythrocytes in autologous plasma with controlled hematocrit of 45% is placed in the setup described in Figure 1. Movie has been initially recorded at 0.2 fps, the rendering at 10 fps is therefore a sped-up of 50 times. The width of the picture is approximately 1 cm (same scale as in Figure 1). The position of the sample is periodically modified by the experimenter to keep the surface of the gel in the field of view.

**Supplemental Movie 2: Sedimentation of control and dyed cells.** (File CellsDyeEffectSpedUp420X.avi) Sedimentation of control (left) and dyed cells (right) in autologous plasma. The initial frame rate was 1 frame per minute. The rendering at 7 fps is therefore a sped-up of 420 times. Both containers are 2 cm high inside and 3mm wide. One can see that the dyed cells have a significantly different sedimentation dynamics than the control ones. Both suspensions have a controlled hematocrit of 45%.

**Supplemental Movie 3: Two-photons microscopy.** (File S0_CombinedSpedUp100Rotated.avi) Sedimentation of a 45% hematocrit suspension of erythrocytes in dyed autologous plasma (with free-acid fluorescein) observed by two-photons microscopy, as illustrated in Figure 2. Recording frame rate is 1 frame every 10 s. The rendering at 10 fps is then 100 times sped-up. The width of the picture is approximately 1.5 mm. The position of the sample is periodically modified by the experimenter to keep the surface of the gel in the field of view.

**Supplemental Movie 4: Light-sheet microscopy.** (File 2021-2-12_3_AnalyzedSpedUp24X.avi) Sedimentation of a suspension of erythrocytes in dyed autologous plasma (Atto647), as observed after reconstruction of surface scan by light-sheet microscopy, as illustrated in Figure 3. Initial hematocrit was adjusted to 45%. A scan of the sample is performed every second, the rendering at 24 fps is therefore 24 times sped-up. The width of the picture is 1.22 mm (same scale as in Figure 3). The movie is recorded lower below the free-plasma/erythrocytes interface, because the position of such interface is hard to detect by the examiner during the scan.
